# Brain structural and genetic correlates of motor coordination and learning behaviours: modelling developmental coordination disorder

**DOI:** 10.64898/2026.06.19.733401

**Authors:** Jason P. Lerch, David Ashbrook, Kamaldeep Gill, Jeffy Rajan Soundara, Jacob Ellegood, John G. Sled, Brian J. Nieman, Jill G. Zwicker, Daniel Goldowitz

## Abstract

Developmental coordination disorder (DCD) is a common neurodevelopmental condition characterized by impaired motor coordination and learning, yet its neurobiological and genetic bases remain poorly understood. Here, we leverage the BXD recombinant inbred mouse panel to model the polygenic architecture of DCD and link behaviour, brain structure, and genotype. High-resolution ex vivo MRI across 14 strains revealed that DCD-like mice have modestly reduced total brain volume, with a distinct neuroanatomical profile characterized by enlarged cortical regions alongside reduced cerebellar, thalamic, and other subcortical volumes. These structural differences closely mirror findings reported in human DCD.

Across strains, variation in brain structure strongly correlated with motor behaviours, with coordinated patterns linking increased cortical and decreased subcortical volumes to poorer motor coordination, while more focal associations were observed for motor learning. Multivariate analysis identified a dominant brain-behaviour axis capturing this cortical-subcortical trade-off. Quantitative trait locus (QTL) mapping revealed multiple loci influencing regional brain volumes, including a prominent locus on chromosome 12 regulating cerebellar structures, but did not identify single loci driving the main multivariate brain-behaviour relationships, consistent with a distributed genetic architecture.

Together, these findings demonstrate that DCD-like motor impairments arise from coordinated alterations across distributed brain systems under polygenic control. This work establishes a translational framework linking genetic variation to brain organization and motor function, and suggests that DCD reflects the extreme of a continuous spectrum of neurobiological variation rather than a discrete condition.

## INTRODUCTION

Developmental coordination disorder (DCD) is a common but under-recognized neurodevelopmental disorder characterized by marked impairments in motor coordination and motor learning that have a significant impact on daily life (Blank et al., 2019). DCD is described in the Diagnostic and Statistical Manual (DSM-5-TR) as having four diagnostic criteria: (1) the acquisition and execution of motor skills are significantly below expected for the person’s age and opportunity for learning; (2) motor skill difficulties significantly interfere with activities of daily living, school performance, prevocational and vocational opportunities, leisure pursuits, and play; (3) onset is in the early developmental period; and (4) the motor difficulties are not better explained by intellectual disability, visual impairment, or another condition affecting movement (e.g., cerebral palsy) (American Psychiatric Association, 2022). DCD often persists into adulthood and has a significant impact on mental health and quality of life throughout the lifespan (Zwicker et al., 2013).

The cause of DCD is unknown but brain differences have been reported. Compared to typically developing children, children with DCD show lower fractional anisotropy and increased axial diffusivity (Brown-Lum et al., 2020), decreased functional connectivity in sensorimotor networks (McLeod et al., 2016, 2014; Rinat et al., 2020), variations in cortical thickness and cortical volume (Langevin et al., 2015; Malik et al., 2024; Reynolds et al., 2017), and smaller gray matter volume in the basal ganglia (Grohs et al., 2021) and several regions of the cerebellum (Gill et al., 2022). Children with DCD also show different patterns of brain activation compared to control children when performing or learning a motor task (Biotteau et al., 2016; Subara-Zukic et al., 2022; Zwicker et al., 2012, 2011).

DCD is also highly heritable (Lichtenstein et al., 2010) but clearly complex - influenced by variation at many locis rather than a single strong variant. This has led to difficulty in confirming the underlying genetic components (Mosca et al., 2016; Mountford et al., 2021; You et al., 2023).

For single gene defects, mouse models have been extremely valuable in illuminating pathogenesis and examining interventions (see, for example Huntington’s Disease (Heng et al., 2008)). Mouse models are less often used to understand complex traits and complex disorders, but has a long history (e.g., (Belknap et al., 1992; Rodriguez et al., 1994; Weimar et al., 1982).The BXD recombinant inbred mice are the largest mouse genetic reference family, and have been used to identify genes underlying complex traits for over 40 years including infectious diseases (Boon et al., 2014), addiction (Reed et al., 2017), and diabetes (Wu et al., 2014). By using this genetically and phenotypically diverse population - reflecting the phenotypic variability seen in patients - we are able to identify novel genes which influence phenotypes of interest and the neuroanatomical mediators of these (Johnson et al., 2025).

To the end of understanding the genetic and structural bases of DCD, developed a model for relating the human phenotype to altered behaviour in the mouse, using the BXD recombinant inbred panel, that links the human phenotype of the disorder to behavioural assays of the mouse. We then examined motor coordination (Rajan et al., 2025) and motor learning (Gill et al., 2023) in 12 strains of BXD mice and their parental strains (C57BL/6J and DBA/2J) and identified five BXD strains that exhibited DCD-like phenotype. DCD-like strains were defined as having problems primarily with gross motor skills, fine motor skills, or with both fine and gross motor skills (Gill et al., 2023)

In the present work, we report the results of imaging the brains of these 14 strains and have identified structural correlates of the behavioural traits that were studied as well as identified genetic loci that modulate both behaviour and neuroanatomy. These findings are then discussed in relation to our current understanding of brain-behaviour-genetic associations of human DCD to provide directions for future research.

## MATERIALS AND METHODS

A total of 193 mice from 14 strains were used in this study, with 101 undergoing behavioural testing plus ex-vivo MRI and the remaining 92 having ex-vivo MRI only. The breakdown of mouse numbers by strain and testing paradigm are given in Table 1.

**Table 1.** Mouse numbers per BXD strain, subdivided by behaviour testing undertaken. Only mice with an MRI scan are included.

| Mouse numbers |  |  |  |  |  |  |  |  |  |  |
| --- | --- | --- | --- | --- | --- | --- | --- | --- | --- | --- |
|  | Overall numbers |  |  | Motor test numbers |  |  |  |  |  |  |
|  | Total | control | motor testing | Motor Learning |  |  |  | Generalized test of motor function |  |  |
|  |  |  |  | Accelerated Rotarod | Complex Wheel | Horizontal Rung | Skilled Reaching | Gait | Open Field | Rotarod |
| BXD1 | 14 | 7 | 7 | 7 | 4 | 7 | 1 | 7 | 7 | 7 |
| BXD15 | 13 | 5 | 8 | 8 | 3 | 8 | 1 | 7 | 7 | 7 |
| BXD27 | 15 | 8 | 7 | 7 | 4 | 7 | 3 | 4 | 4 | 4 |
| BXD28 | 18 | 10 | 8 | 8 | 4 | 6 | 4 | 7 | 7 | 7 |
| BXD32 | 15 | 9 | 6 | 6 | 3 | 6 | 3 | 5 | 5 | 5 |
| BXD40 | 13 | 7 | 6 | 6 | 3 | 6 | 2 | 5 | 5 | 5 |
| BXD45 | 16 | 8 | 8 | 8 | 4 | 8 | 4 | 4 | 4 | 4 |
| BXD65a | 16 | 8 | 8 | 8 | 3 | 8 | 4 | 8 | 8 | 8 |
| BXD69 | 5 | — | 5 | 5 | 2 | 3 | 3 | 5 | 5 | 5 |
| BXD75 | 15 | 7 | 8 | 8 | 2 | 7 | 4 | 5 | 5 | 5 |
| BXD81 | 11 | 2 | 9 | 7 | 3 | 8 | 4 | 8 | 8 | 8 |
| BXD86 | 14 | 7 | 7 | 7 | 2 | 7 | 4 | 4 | 4 | 4 |
| C57BL/6J | 15 | 7 | 8 | 8 | 2 | 4 | 4 | 8 | 8 | 8 |
| DBA/2J | 13 | 7 | 6 | 6 | 2 | 3 | 4 | 6 | 6 | 6 |
| Total | 193 | 92 | 101 | 99 | 41 | 88 | 45 | 83 | 83 | 83 |

### Behaviour testing

The behavioural tests that were carried out were those described in previous published work to assess motor coordination (Rajan et al., 2025) and motor learning behaviour (Gill et al., 2023) in this cohort of BXD strains. For motor co-ordination, we assessed gait, open field, and rotorod behaviours. For motor learning, we assessed performance on an accelerated rotorod, a complex wheel, a horizontal rung, and skilled reaching task. The details of these tasks can be found in the citations noted above with the numbers used in the present study shown in Table 1.

### MRI

Mice were anesthetized and perfused at ∼1mL/min via the left ventricle with 10ml of 0.1M PBS containing 10U/mL heparin and 2mM ProHance (a Gadolinium contrast agent) followed by 30mL of 4% paraformaldehyde (PFA) containing 2mM ProHance at the same perfusion rate. After perfusion and initial fixation, mice are decapitated; the brain and skull maintained in 4%PFA + 2mM ProHance overnight at 4°C before being transferred to 0.1M PBS containing 2mM ProHance and 0.02% sodium azide for at least 7 days prior to the dissection of the head with brain minus skin, jaw, and eyes.

MRI scans were conducted on a multi-channel 7.0 Tesla MRI scanner (Agilent Inc., USA) at the Mouse Imaging Centre at the Hospital for Sick Children in Toronto. Sixteen custom-built solenoid coils enabled imaging the brains in parallel (Nieman et al., 2018) using the following parameters for the anatomical MRI scans: T2-weighted, 3-D fast spin-echo sequence, with a cylindrical acquisition of k-space (Spencer Noakes et al., 2017) and with a TR of 350 ms, and TEs of 12 ms per echo for 6 echoes, field-of-view of 20 x 20 x 25 mm3 and matrix size = 504 x 504 x 630 giving an image with 0.040 mm isotropic voxels. Total imaging time was ∼14 h.

To visualize and compare any changes in mouse brain anatomy, images were segmented using the MAGeT algorithm (Chakravarty et al., 2013) into 590 different brain areas (Beera et al., 2018; Dorr et al., 2008; Qiu et al., 2018; Richards et al., 2011; Steadman et al., 2014; Ullmann et al., 2013). Statistical analyses were then carried out on these brain volumes, correlating them with behavioural outcomes or group status. Multiple comparisons were controlled using the False Discovery Rate (FDR) (Benjamini and Hochberg, 1995). Canonical Correlation Analyses were implemented using the *mixOmics* package in R (Rohart et al., 2017) using its built in shrinkage algorithm for regularization.

### QTL mapping

For each brain region, strain mean volumes were calculated for 1) all mice, 2) mice who had received no behavioural phenotyping (no motor training), 3) mice who had received behavioural phenotyping (motor training). This allowed us to build confidence in our QTLs as (2) and (3) are independent groups, and so finding the same QTL region in both of genes groups increases our confidence in that QTL. Quantitative trait locus (QTL) mapping was then carried out using R/qtl2 (Broman et al., 2019). The mean value per strain was calculated to reduce environmental variations, increasing the ability to detect QTLs. The BXD family has been produced in several “epochs,” using both standard F2 recombinant inbred methods and advanced intercross recombinant inbred methods (David G. Ashbrook et al., 2021), leading to both expected and unexpected kinship between BXD types that can introduce bias. Therefore, a kinship matrix was used in R/qtl2. Genotypes for 21,056 markers derived from whole genome sequencing (WGS) of the BXD family were downloaded from GeneNetwork.org (Sloan et al., 2016; Villani et al., 2025). A −logP of > 4 was defined as nominally genome-wide significant, and >3 as suggestive. To identify genes within the QTL, a 1.5 −logP drop interval was applied around the peak position, providing approximately a 95% confidence interval (Manichaikul et al., 2006).

### Use of Artificial Intelligence Generated Content

Artificial intelligence tools such as LLMs were not used in the preparation of this manuscript

## RESULTS

We separated the 14 strains (12 BXD crosses plus C57Bl6/J and DBA/2, the two founder lines) in our study into 5 DCD-like and 9 non-DCD-like based on their behaviour on a battery of motor co-ordination (Rajan et al., 2025) and motor learning tasks (Gill et al., 2023). Half the mice from each strain were tested on 7 motor coordination and motor learning assays, the other half were naive. All mice were perfusion fixed and the brains, left within the skull, underwent high resolution structural MRI imaging followed by automatic segmentation into 590 brain areas (see Methods). We first tested whether the anatomy of the 5 DCD-like BXD strains differed from the 9 non-DCD-like strains. Overall, as seen in Figure 1, DCD-like mice had 2.6% smaller brain volumes (t=-2.6, p=0.014). After controlling for overall brain volume, DCD-like mice featured larger cerebral cortices and smaller subcortical volumes (Figure 1C). Notable increases included cingulate cortex area 24b and 29c, the orbital area, parietal association cortex, primary somatosensory cortex, secondary motor area, and primary motor cortex, along with the corpus callosum and cingulum bundles. Notable decreases were found in the thalamus, cerebellar nuclei, stria terminalis and stria medullaris, and the mammilothalamic tract. These volume differences correlated with motor coordination and learning behaviours, as shown for two illustrative examples in Figure 1D-E. These results indicate that brain structure differs between DCD-like and non-DCD-like mice and support the concept that neuroanatomy relates to motor coordination and learning.

**Figure 1:**
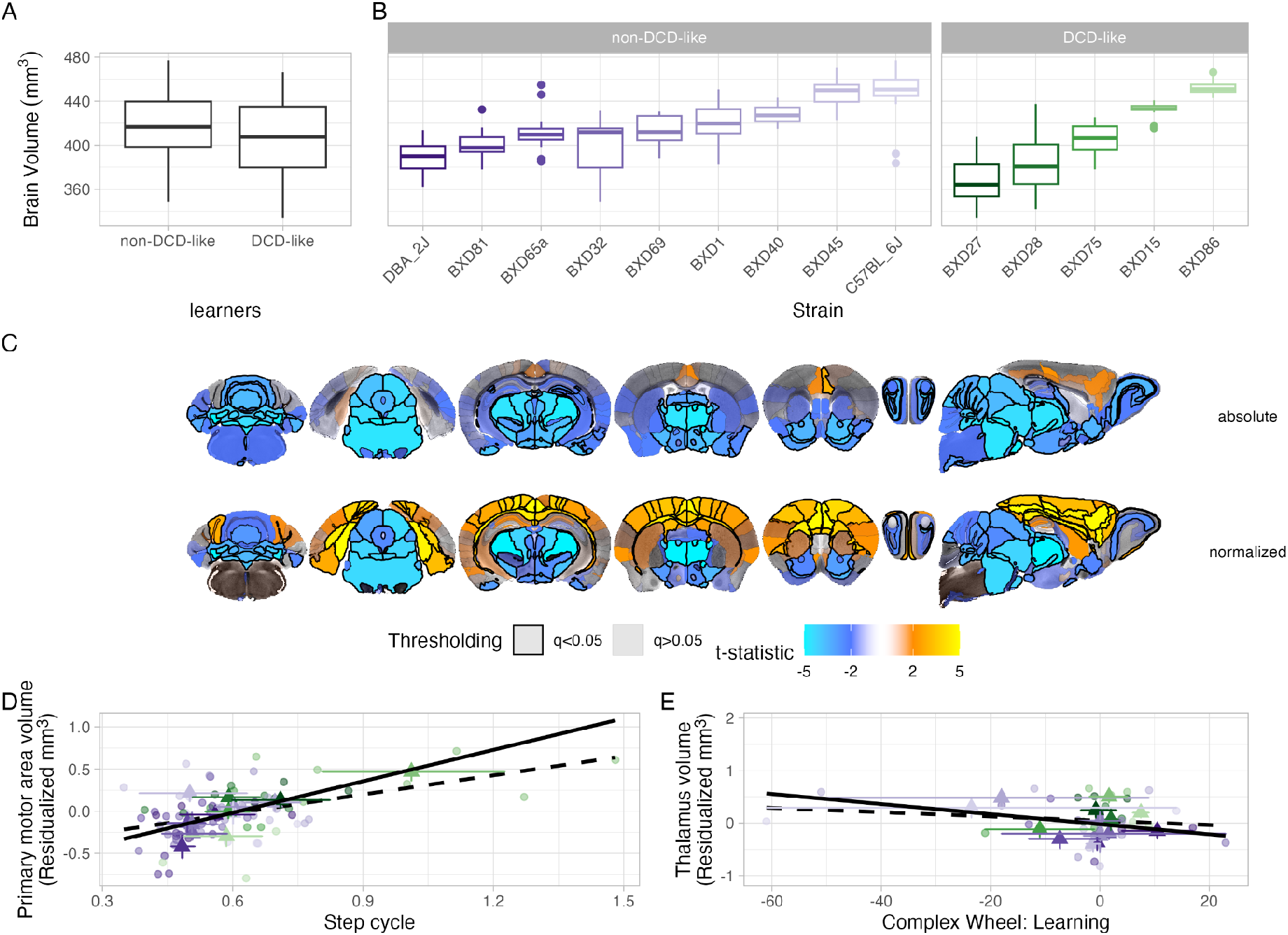
Comparison of DCD-like vs non-DCD-like mouse strains. DCD-like strains showed slightly smaller overall brain volume, as seen in (A) and separated by strain in (B). DCD-like mice have larger cerebral cortices and smaller subcortical volumes, which is especially pronounced when normalizing for overall brain volume (C). The volumes are associated with coordination and motor learning behaviours, with the primary motor area correlating positively with step cycle (D) and thalamus volume correlating negatively with complex wheel learning (E). Least-squares fits are shown both across strains (solid line) and across mice (dashed line).

To test these brain-behaviour associations between neuroanatomy and motor coordination and motor learning behaviours, we employed a series of univariate tests. Given the stronger associations between brain and behaviour when comparing means of our 14 strains than for pooling the individual mice, we computed mean volumes per strain for each of the 590 brain regions in our hierarchical atlas and mean scores per strain for each of the 25 behavioural scores across 7 behavioural assays in our battery. Pearson correlations for each brain region, after controlling for overall brain volume, and for each behavioural score are shown in Figure 2. Multiple large correlations were identified, at times exceeding |r|>0.9.

**Figure 2:**
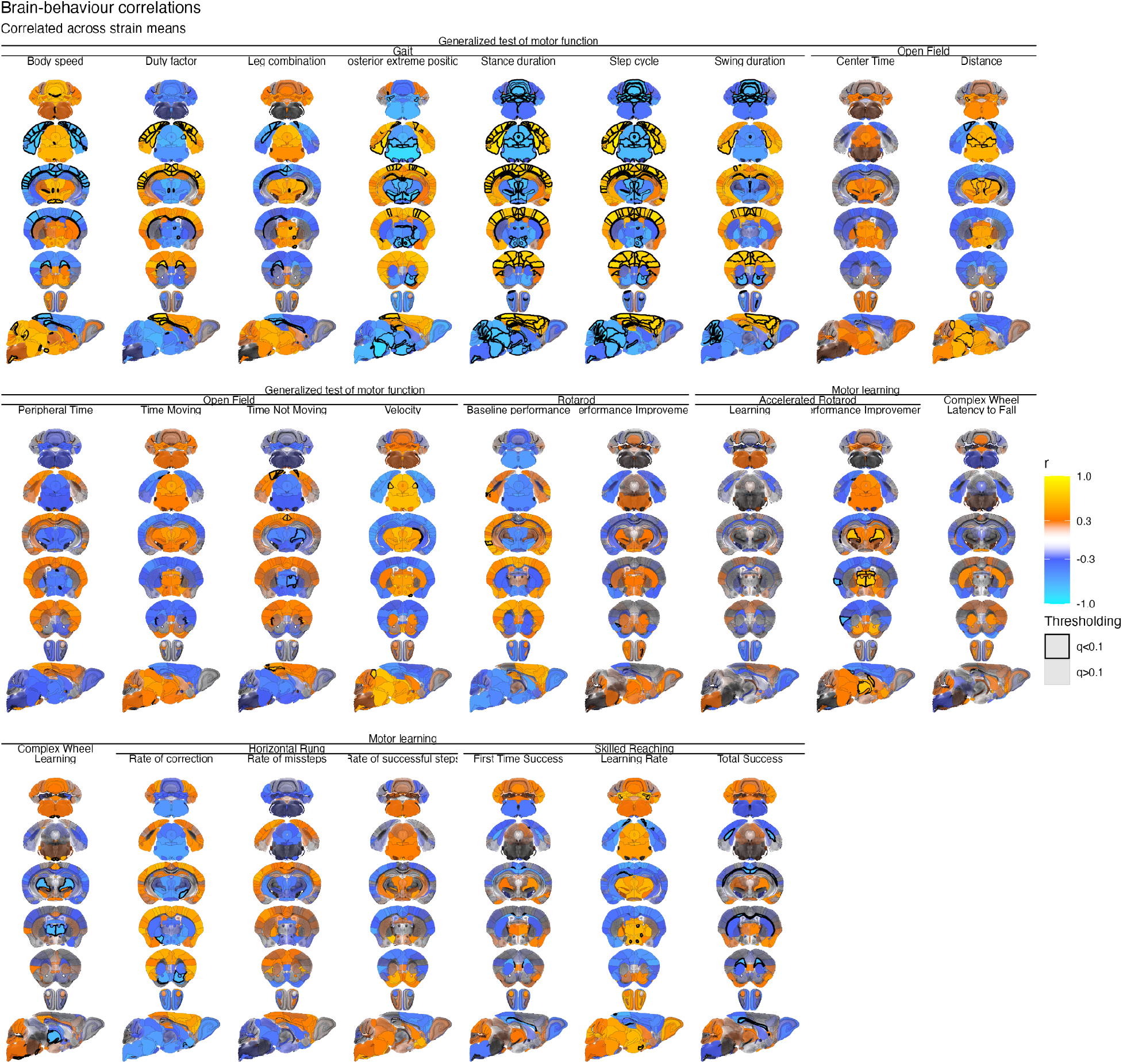
Brain-behaviour correlations. Mean volumes per strain were computed for every brain region and correlated against mean behaviour scores. Orange-red colours indicate that as that behaviour score increases, so does brain volume, whereas blue colours indicate that behaviour score increases are associated with brain volume decreases. Brain volumes were adjusted for overall brain volume.

To investigate these brain-behaviour relations in more detail,we restricted ourselves to key motor brain areas and plotted them as a clustered heatmap in Figure 3. Key correlations identified were:

**Figure 3:**
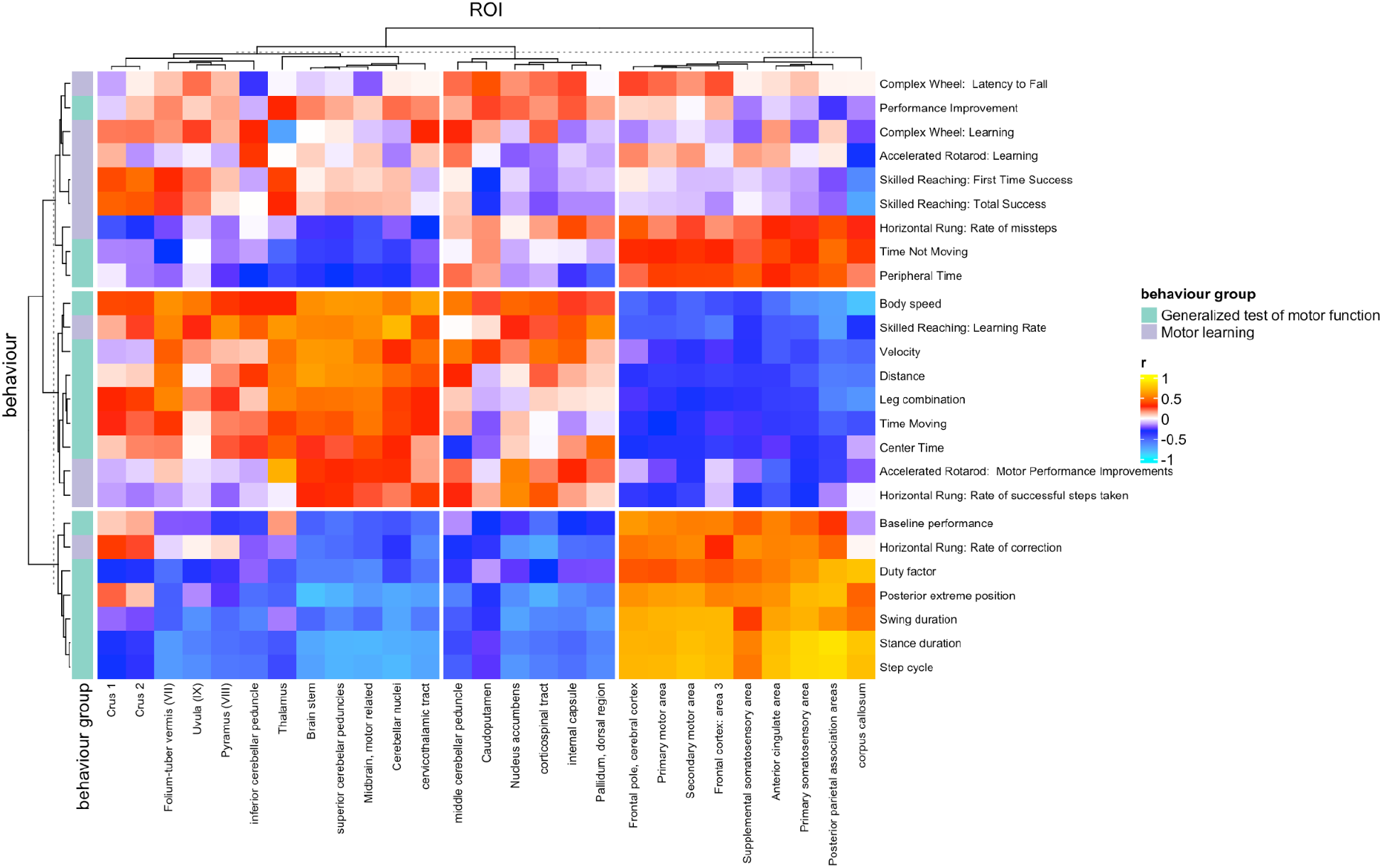
Brain-behaviour correlations in key motor areas of the brain. We selected key nodes in the motor network and plotted their correlation (red/yellow = positive correlation, blue/aqua = negative correlation) against all 25 scores from our 7 motor coordination and motor learning tasks. The heatmap was then clustered along both axes. The clustered heatmap was divided into 3 dominant patterns on each axis to aid in visualization and interpretation.

1. higher gait scores (step cycle, stance duration, duty factor, posterior extreme position) associated with
  a. larger cortical motor regions (primary and secondary motor areas and key association areas) and the corpus callosum
  b. smaller cerebellar and subcortical (pallidum, thalamus) sizes.
2. higher body speed, velocity and distance in the open field, and skilled reaching learning rate associated with
  a. larger cerebellar nuclei, brain stem, thalamus, cerebellar peduncles and cortico-thalamic tract
  b. smaller corpus callosum, parietal association area, and, to a lesser extent, smaller remaining cortical motor regions

Motor learning tasks were more focally associated with brain regions than the broader patterns identified with motor coordination behaviours. Skilled reaching was associated with smaller corpus callosum volumes, and complex wheel learning and accelerated rotarod improvements with smaller thalamus volumes.

Finally, we tested for multivariate brain-behaviour relations using canonical correlation analyses (CCA). As shown in Figure 4, 8 components retained a correlation of r ≥ 0.5, with the first component capturing the dominant pattern of larger cortices alongside smaller cerebellar and subcortical regions positively associated with step cycle, stance duration, swing duration, posterior extreme position, and duty factor and inversely associated with body speed, velocity, and skilled reaching learning rate.

**Figure 4:**
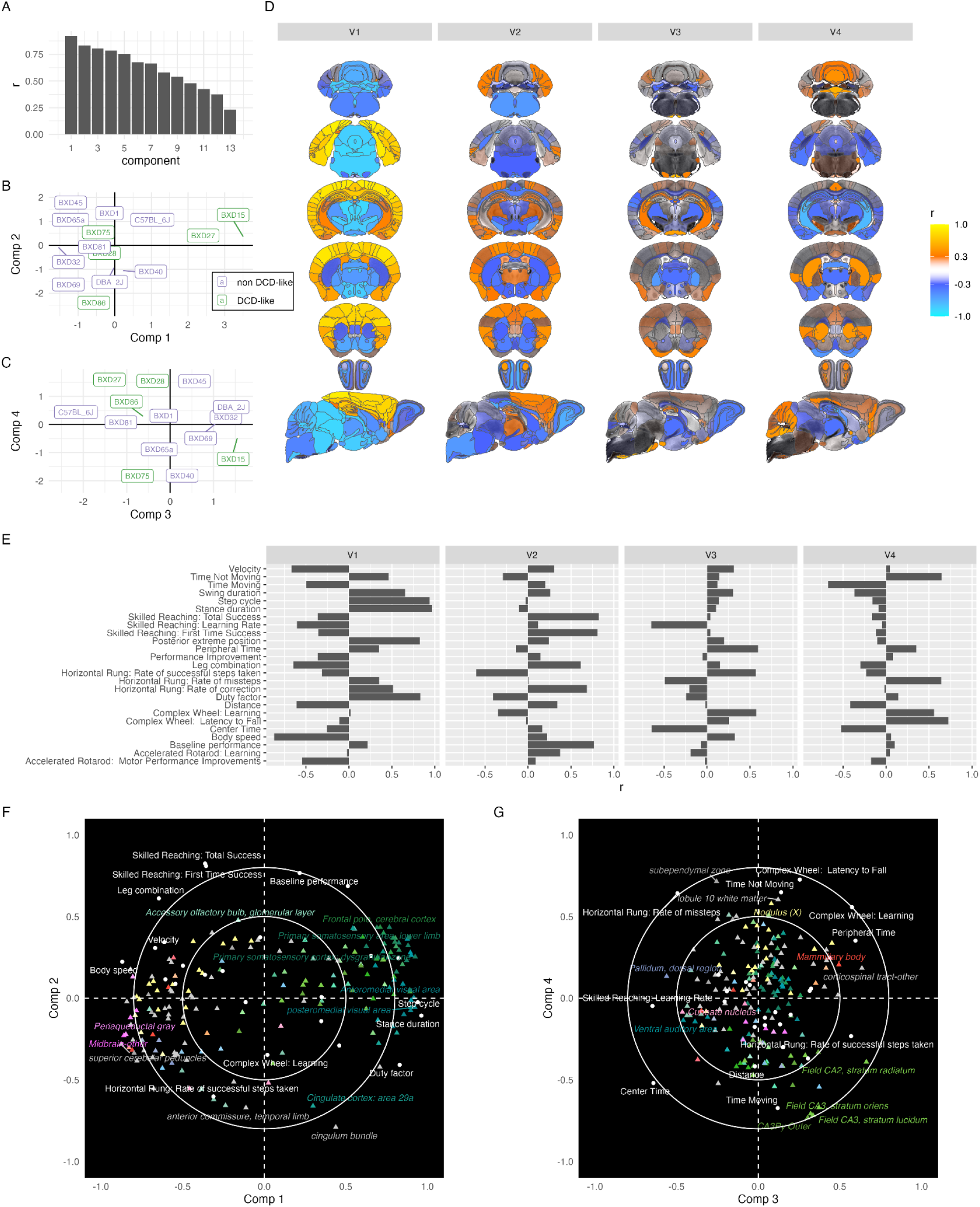
Sparse canonical correlation analyses show multivariate relations between brain and behaviour. As shown in (A), 8 components retained a correlation greater than 0.5. (B-C) show the distribution of the 14 strains across the first four components; the DCD-like and non-DCD-like strains are interspersed amongst each other, with only the first component showing statistically significant separation (p=0.017) between groups. (D) illustrates how the first four components load onto the brain, (E) how they load onto behaviour, and (F-G) the biplots of brain-behaviour relations (with colours taken from the Allen atlas).

## QTL analyses

We next sought to understand the genetics underlying these brain behaviour associations. For the volume of each brain region, we carried out QTL analyses on strain means of 1) all mice, 2) mice who did not receive any motor phenotyping, and 3) strain means of mice who did receive motor phenotyping. This produced 201 nominally suggestive (LOD > 3.5) and 92 nominally significant (LOD > 4) QTL [supp table]. We detected several clusters of QTL - genomic regions that appear to regulate the volume of several related brain regions. One of these appears particularly interesting, as it controls cerebellar volume.

This QTL is on chr12 110.7-120 Mb, and influences the volume of the Ansiform lobule, crus 1 and 2, crus 1 and 2 white matter, hemispheric regions, paramedian lobule, and the simple lobule (Table 1). The fact that several of these are significant in both the motor and no motor groups gives us even greater confidence, as these are two separate sets of individuals. We additionally ran QTL analyses on the top components of the brain-behaviour CCA. None of the first three components were associated with any QTLs, though components 4 and 8 had QTLs that were nominally significant (V4 LOD = 4, chr7:59.99-74.52 Mb ; V8 LOD = 6.1 chr8:121.59-125.16 Mb). This suggests that the main genetic associations are with individual components of the DCD network rather than the DCD network as an entity.

**Table 1.** Anatomical regions sharing a QTL on chromosome 12. Region names, QTL locations, and the LOD score are given for each.

| *Brain region* | *Chromosome* | *Position (Mb)* | *Max LOD* | *Confidence interval* |
| --- | --- | --- | --- | --- |
| *Ansiform lobule, all samples* | 12 | 113.82 | 6.59 | 113.81-119.92 |
| *Ansiform lobule, samples with motor tasks* | 12 | 113.82 | 6.47 | 113.81-119.92 |
| *Ansiform lobule, samples without motor tasks* | 12 | 113.82 | 5.51 | 113.81-119.92 |
| *crus 1 white matter, all samples* | 12 | 114.86 | 3.42 | 113.81-119.92 |
| *Crus 1, all samples* | 12 | 115.06 | 4.28 | 113.81-119.92 |
| *Crus 1, samples without motor tasks* | 12 | 114.71 | 4.47 | 113.81-119.92 |
| *crus 2 white matter, all samples* | 12 | 113.82 | 3.96 | 112.51-119.92 |
| *crus 2 white matter, samples with motor tasks* | 12 | 114.86 | 3.14 | 113.81-119.92 |
| *crus 2 white matter, samples without motor tasks* | 12 | 113.82 | 3.73 | 112.51-119.92 |
| *Crus 2, all samples* | 12 | 113.82 | 6.35 | 113.81-119.92 |
| *Crus 2, samples with motor tasks* | 12 | 113.82 | 6.74 | 113.81-119.92 |
| *Crus 2, samples without motor tasks* | 12 | 113.82 | 5.13 | 112.51-119.92 |
| *Hemispheric regions, samples with motor tasks* | 12 | 113.82 | 3.51 | 112.51-119.92 |
| *Paramedian lobule, all samples* | 12 | 113.82 | 3.54 | 110.75-119.92 |
| *Paramedian lobule, samples with motor tasks* | 12 | 113.82 | 4.21 | 113.81-119.92 |
| *Simple lobule, all samples* | 12 | 115.07 | 3.12 | 113.81-119.92 |
| *Simple lobule, samples without motor tasks* | 12 | 115.07 | 3.68 | 113.81-119.92 |
| *trunk of crus 2 and paramedian white matter, samples with motor tasks* | 12 | 115.38 | 3.18 | 112.51-119.92 |

## Discussion

We investigated brain phenotypes and brain-behaviour relations in a polygenic mouse model of DCD. We found that the DCD-like mice had altered neuroanatomy compared to non-DCD-like mice, including having slightly smaller brains, prominent reductions in the cerebellum and subcortical and brain-stem regions alongside increases in volume in the neocortex. These patterns largely match human brain imaging findings, especially in the cerebellum, thalamus, and basal ganglia. (Gill et al., 2022) showed that children with DCD have smaller cerebellar volume in several regions, including the right and left crus I, right crus II, left VI, right VIIb, and right VIIIa lobules, compared to typically developing children. Lower motor performance was related to smaller cerebellar grey matter volume. Compared to typically developing children, children with DCD have lower mean volume in the left and right pallidum (part of the basal ganglia) but these findings were not related to motor performance (Grohs et al., 2021). Very preterm born children with/at-risk of DCD also show smaller brain volume in the pallidum and thalamus (Dewey et al., 2019).

Findings in the neocortex in humans are more mixed. (Malik et al., 2024) showed that relative to neurotypical children, children with DCD had significantly greater grey matter volume in the left superior frontal gyrus, which correlated with poorer motor performance. They hypothesized that children with DCD may have delayed or absent cortical thinning, potentially due lack of pruning as seen in other neurodevelopmental disorders (e.g. congenital amusia (Hyde et al., 2007) or autism (Tang et al., 2014)). In contrast, (Langevin et al., 2015) reported thinner cortex in the right temporal pole and (Reynolds et al., 2017) found smaller grey matter volume in the right middle, medial, and superior frontal gyri in children with DCD. Small samples alongside subtle differences in acquisition and data processing methods and participant ages likely account for these discrepancies.

We next tested for how variations in the brains of our mice reflect variations in motor coordination and motor learning. The dominant brain pattern again involved increases in the neocortex alongside decreases in the cerebellum, brain-stem, and subcortical regions. This pattern was associated primarily with generalized motor function, with more focal brain changes associated with variations in motor learning. DCD-like mice only partly separated from non-DCD-like mice, suggesting that the brain alterations that determine developmental coordination exist on a continuum with DCD at one end, in agreement with previous theories (Kaplan et al., 2006, 1998; Pearsall-Jones et al., 2010).

Our QTL analyses provide an intriguing window onto the complex genetic architecture of DCD. We identified multiple QTLs for brain volumes of those regions separating DCD-like from non-DCD-like mice. Yet we could identify no QTLs for the first multivariate components (Figure 4) that associate motor behaviours with brain phenotypes (though we did find QTLs for components 4 and 8). This suggests that separate genetic variants influence the brain circuitry that ultimately governs motor coordination, and that multiple variants are needed to tip into DCD. Yet there are intriguing links of our findings to monogenic disorders. 22q11.2 deletion, for example, features comorbid motor difficulties (Cunningham et al., 2018), and previous mouse brain imaging studies have shown a similar pattern of decreases in cerebellar and concomitant increases in neocortical volumes (Ellegood et al., 2014). Multiple monogenic developmental disorders have associated motor coordination phenotypes, yet not all feature a DCD-like brain phenotype. We have recently looked across a broad swath of autism-related mouse models, and found one cluster of these models that features a brain pattern similar to what we see in DCD; these models are preferentially enriched for nervous system development genes, in particular acting on the WNT and MAPK signalling pathways (Ellegood et al., 2025). In summary, we argue that common variant DCD, as modelled in this manuscript, requires the action of distinct genes acting on distinct brain regions, yet that these brain patterns are similar to what can be found in single gene mutations or CNVs (such as those found in human DCD (Mosca et al., 2016)).

The very high brain-behaviour correlations we identified are worthy of additional comment. Human brain imaging studies investigating relations between brain structure or function and cognition or behaviour tend to find low correlations coefficients, usually |r| < 0.3, needing thousands of participants to be reliable (Marek et al., 2022). Here we identified high brain-behaviour correlation coefficients, exceeding |r| > 0.9 in several cases. It is important to note that correlating across individuals results in |r| maxing out at about 0.5; this is still higher than usually found in human studies, but less than correlating across strains. We draw three conclusions relating to brain-behaviour correlations from these data. First, our results suggest that behaviour and brain structure are tightly coupled, with the low correlations identified in human studies attributable to large amounts of statistical noise. Second, we can point to two likely sources of noise: (1) that correlations across individual mice are higher than across individual humans suggests that the complex environment and experience of humans is inherently noisy, and (2) that correlations across strains are much higher than across individual mice suggests that averaging out noise and/or idiosyncratic plasticity on both the brain and behaviour side significantly increases brain-behaviour associations.

There are several caveats to note. Most importantly, we only used 14 strains, thus significantly limiting our statistical power to detect QTLs and increasing the probability of false positives, and those that were detected were too large to reliably pinpoint individual genes or variants. We also relied on MRI as our readout of brain architecture; while it has the significant advantages of providing unbiased whole brain coverage and being analogous to measures acquired in human cohort studies, MRI-derived volumes do not easily yield cellular or molecular explanations. Finally, brain imaging and behavioural testing were conducted in young adulthood, thus eliding developmental explanations for DCD.

## Acknowledgements

This work was funded by a Brain, Behavior and Development Catalyst Grant (co-PIs Goldowitz and Zwicker), BrainCanada and a CIHR Platform grant (PI Lerch), and a Trainee Boost Award (Soundara Rajan) from the BC Children’s Hospital Research Institute (BCCHRI). Gill was funded by the Effie I. Lefeaux Scholarship, Syd Vernon Graduate Student Award, and Cordula and Gunter Paetzold Fellowship. Zwicker is funded by BCCHRI, Canadian Institutes of Health Research, and Canada Research Chairs Program. For the purpose of open access, the author has applied a CC BY public copyright licence to any Author Accepted Manuscript version arising from this submission.

## Code availability

Code for all analyses in this manuscript can be found at https://github.com/jasonlerch/BXD-DCD-MRI

## Notes

### Competing Interest Statement

The authors have declared no competing interest.

### Summary of Updates

Reformatted for Genes, Brain and Behavior

